# Specialization of male reproductive tactics correspond with large gonads and small brains

**DOI:** 10.1101/2025.03.22.644732

**Authors:** Helen Stec, Grace Y. Zhang, Ben A. Sandkam

## Abstract

Sexual selection has strong effects on gonad size, which has been proposed to shift energetic allocations resulting in concomitant decreases to brain size. However, mixed findings leave it unclear whether negative correlations arise from direct energetic trade-offs or correlated selection. We tested whether male reproductive tactics impose energetic trade-offs by comparing brain and gonad sizes in *Poecilia parae*, a fish with discrete alternative male morphs specializing in three reproductive strategies: coercion, display, and sneaking. The obligate sneaker morph had substantially larger gonads and smaller brains than the other morphs, consistent with an energetic trade-off. However, examining individuals within morphs revealed a positive relationship, contradicting energetic trade-offs. To resolve which morphs reflect the ancestral state, we examined two closely related species whose males utilize more flexible reproductive strategies, *Poecilia picta* and *Poecilia reticulata*. Within these species, a negative correlation between gonad and brain size was observed, consistent with correlated selection shaping traits towards multiple reproductive peaks. Additionally, neuron-to-glia ratio (a proxy for energetic demands) showed no link to gonad size. Our results suggest that reproductive strategies shape brain evolution through correlated selection rather than direct energetic trade-offs, challenging assumptions of sexually selected traits imposing constraints through direct resource allocation.

## Introduction

Sexual selection is a powerful evolutionary force acting on traits and tissues that are energetically costly to produce and maintain (Darwin, 1871; Rowe & Rundle, 2021). This high cost leads to a tight co-evolution between mating strategies and the size of relevant traits. A classic example of this evolutionary response is the positive correlation between male gonad size and the prevalence of promiscuity in the mating strategy across species (Kenagy & Trombulak, 1986; Pitnick et al., 2001). Such selection to increase gonad size has been proposed to reduce brain size due to trade-offs between energetically costly tissues, in line with the ‘Expensive Tissue Hypothesis’ (Aiello & Wheeler, 1995; Heldstab et al., 2022; Lemaître et al., 2009). However, comparative studies have revealed mixed support for this dynamic. While some have confirmed such a negative relationship between brain and gonad size (bats; Pitnick et al., 2005; swordtail fish; Stec et al., 2023), others have found a positive relationship (odontocetes; Kelley et al., 2014; frogs; Jin et al., 2015, mammals; Husak et al., 2024). Such conflicting support calls for a more focused evaluation of within and across species variation to determine if the evolution of mating strategies either influences brain evolution through direct trade-offs of energetic allocation with gonad size or if there is co-selection on these two traits.

Two potential causes have been proposed to explain the discrepancies in support of brain evolution through energetic trade-offs: (1) confounds with ecological niche, (2) energetic trade-offs at the cellular level. Most tests of the expensive tissue hypothesis have been comparative studies examining diverse species that also differ dramatically in ecological niche. These differences in ecology often correspond to trade-offs in brain and gonad size (Pitnick et al., 2005), prompting some to caution the interpretation of gonad/brain trade-offs as they are confounded with total energy available (Gonda et al., 2013; Heldstab et al., 2022), and predicting evolutionary trade-offs is difficult when both acquisition and allocation are allowed to vary (Roff & Fairbairn, 2007; Van Noordwijk & De Jong, 1986). Therefore, testing if energetic trade-offs with gonad size influence brain evolution would require systems with little to no difference in ecological niche.

A second possibility for the mixed support of the expensive tissue hypothesis is that the principles of energetic trade-offs shape brain evolution through alterations to cellular composition rather than raw brain size. The brain is primarily comprised of two cell types: neurons and glial cells, with energetic demands of neurons far surpassing those of the glia (Attwell & Laughlin, 2001; Harris et al., 2012). Therefore, energetic trade-offs between tissues could be accomplished without altering brain size, but instead by shifting the ratio of neurons to glia. The neuron to glia ratio has long been known to differ across species (Herculano-Houzel, 2014; Reichenbach, 1989) and even differs across closely related species (Herculano-Houzel, 2014), indicating this trait may be highly amenable to selection. Therefore, testing whether energetic trade-offs with gonad size shape brain evolution also requires testing for shifts in the cellular composition of the brain.

Brain size evolution through the Expensive Tissue Hypothesis is a specific case of the acquisition-allocation model (the ‘Y-model’) for the evolution of two traits under a trade-off. This model suggests that the evolution of two traits is constrained due to having a finite resource and allocating more resources to one trait necessitate a decrease in resources to the other trait (De Jong & Van Noordwijk, 1992; Garland et al., 2022; Roff & Fairbairn, 2007). The Y-model of trait evolution provides two clear expectations to test whether reproductive tactics shape brain evolution through energetic trade-offs with gonad size: increases in gonad size will be accompanied by concomitant decreases to brain energetic demands both (a) between reproductive tactics, and (b) within reproductive tactics (Roff et al., 2002; Roff & Fairbairn, 2012). Alternatively, if negative correlations between brain and gonad sizes are not due to strict energetic trade-offs but instead are the result of brains and gonads being under co-selection, then increases to gonad size will be accompanied by decreases to brain energetic demands between reproductive tactics, but not within reproductive tactics (Agrawal, 2020).

Systems with genetically determined alternative reproductive tactics provide a powerful way to test whether mating strategies shape brain evolution through trade-offs with male gonad size. Males of the freshwater fish *Poecilia parae* always occur as one of five distinct genetically-determined morphs that are inherited through the Y chromosome (Lindholm et al., 2004; Sandkam et al., 2021). All five morphs co-occur within populations and occupy the same ecological niche (Liley, 1966; Lindholm et al., 2004). Despite sharing ecological niche, morphs differ dramatically in their mating strategies, specializing either in display, coercion, or sneaker strategies (Hurtado-Gonzales & Uy, 2009; Lindholm et al., 2004). The ‘display’ strategy is used by all three of the ‘melanzona’ morphs, which attract females through colourful courtship displays of their horizontal stripe (red, yellow, or blue). The ‘coercion’ strategy is used by the ‘parae’ morph, which has a larger body and vertical black stripes, which it uses to chase away competitors and aggressively force-copulate with females. Meanwhile, the ‘sneaker’ strategy is used by the ‘immaculata’ morph, which resemble juvenile females and sneak up to force-copulate with females.

Males that specialize in sneaker reproductive strategies experience strong selection to increase gonad size (Gross, 1996; Mank, 2023). As predicted, the gonads of the immaculata morph of *P. parae* have been observed to be larger than those of the other morphs (Hurtado-Gonzales & Uy, 2009). This dramatic difference in gonad size between genetically determined alternative mating strategies presents an opportune system to test whether mating strategies shape brain evolution through energetic trade-offs with gonad size or if traits are being co-selected, all while avoiding confounds of differences in ecological niche. If energetic trade-offs shape brain evolution, then the immaculata morph should also have smaller brains than the other morphs, and this negative relationship should also be observed among individuals within each morph.

Since the specialized morphs of *P. parae* are all in the same species, a straight comparison across morphs would not clarify whether selection has acted to increase gonad size in the immaculata morph or decrease gonad size in the melanzona and parae morphs. The species *Poecilia picta* and *Poecilia reticulata* provide a point of comparison to resolve which morphs are under selection as they can act as a proxy for the ancestral state (Lindholm et al., 2015). Not only do these species occur in sympatry, but they are close relatives (Fig. 1A; 15.5 and 18.8 million years ago, respectively; Rabosky et al., 2018) and males utilize a flexible mating strategy. Rather than specializing in coercion, displaying, or sneaking, males *of P. picta* and *P. reticulata* will use all three strategies depending on social context (Houde, 1997; Liley, 1966; Pilastro & Bisazza, 1999). By comparing brain/gonad allocation in the specialized morphs of *P. parae* to their sympatric sister taxa, we can infer if selection has acted to increase male gonad size in the immaculata morph and test for potential trade-offs with the brain.

**Figure 1.**
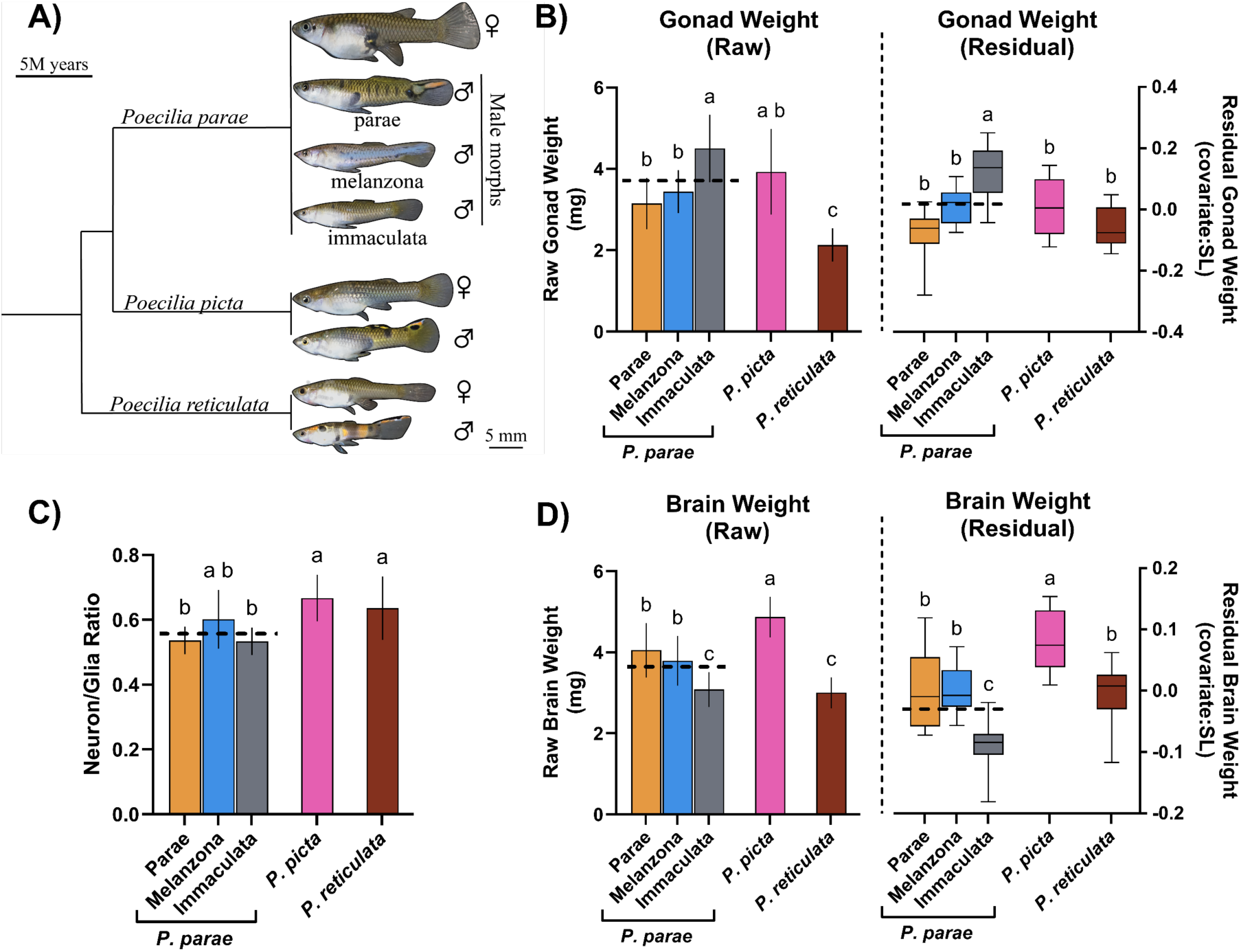
The male morphs of *P. parae* (parae, melanzona, and immaculata) specialize in different mating strategies (coercion, display, and sneak), while males of the closely related species *P. picta* and *P. reticulata* flexibly utilize mating strategies (A; diverged ∼15.5 and ∼18.8 MYA, respectively; Rabosky et al., 2018). The immaculata morph of *P. parae* (an obligate sneaker) has the largest testes (B), both in raw weight (left), and when correcting for body size (right). Meanwhile, the brains of the immaculata morph are the smallest (D), both in raw weight (left) and when correcting for body size (right). Within the brain, both the parae and immaculata morphs had a lower neuron/glia ratio compared to the sister species (C). Horizontal dashed lines indicate the mean across morphs for *P. parae.* Two-way ANOVAs followed by Tukey comparisons were used. Error bars indicate SE. Significant differences between groups (p< 0.05) are indicated by different letters.

In this study, we compared gonad size and brain traits to resolve the two alternative hypotheses for explaining the negative relationships sometimes observed between gonad size and brain size: (1) If male reproductive tactics drive selection on brains through energetic trade-offs with gonad size, then larger male gonads should always correspond to brains that are less energetically costly (i.e. are smaller or have a lower neuron/glial ratio) both across and within the morphs of *P. parae* and their close relatives. Alternatively, (2) if male reproductive tactics drive correlated selection on gonads and brains, then larger male gonads will correspond to less energetically costly brains across morphs but not within morphs.

## Materials and Methods

### Animal Model

Laboratory stocks of *P. parae, P. reticulata,* and *P. picta* were established from wild-caught sympatric populations from Suriname in June 2022. To avoid any potential differences due to developmental plasticity, all individuals were raised in a common garden design where all aquaria were part of the same recirculating filtration system (∼4,500 l total) and each tank contained gravel, artificial plants for cover, and algae. Full-spectrum lighting was provided by LED lamps directly above each tank and kept on a 12:12h light:dark cycle. Water temperature was maintained between 25-27 °C. Fish were fed *ad libitum* twice a day on a mixed diet of live brine shrimp and pellets (Fancy Guppy, Kyorin Co., Himeji, Japan). All individuals used in this study were sexually mature, in Poeciliid fishes males stop growth upon reaching sexual maturity, therefore any variation in tissue size was not due to age (Klaus D. Kallman, 1983; Reznick & Endler, 1982).

For *P. parae,* ten females and ten males were sampled for each of the three major morphs: parae, immaculata, and melanzona. To keep sample sizes of each reproductive strategy the same we only examined one of the three melanzona morphs (blue melanzona) since all three utilize the same reproductive strategy (Hurtado-Gonzales & Uy, 2009) and are highly similar at the genetic level (Sandkam et al., 2021). For *P. reticulata* and *P. picta* ten males and ten females were used for each species.

### Dissections

To compare investment differences between brains and gonads, we developed a precise dissection protocol that ensured standardization and allowed for accurate group comparisons. Individuals were sacrificed using MS-222, dried with a Kimwipe (Kimtech, Roswell, Georgia), and weighed using a Pioneer Precision scale (Ohaus, Parsippany, New Jersey, USA) before making standardized photographs on a dissection scope that included a ruler (Leica Microsystems, Wetzlar, Germany). Images were analysed using FIJI to determine body size (standard length, SL, distance from snout to end of caudal peduncle) (Schneider et al., 2012). Whole brains (including olfactory bulbs) were removed by cutting rostrally at the second vertebrate in contact with the first plural rib (<u>Fig. S1</u>) and cutting optic nerves at the optic chiasm. Dissecting brains using precise landmarks is key to ensuring weight differences are not due to inconsistent dissections. Female gonad weights were not examined since females of these species are livebearers, and their gonad weight fluctuates dramatically across their pregnancy cycle (Thibault & Schultz, 1978). To prevent tissues from drying out and giving inaccurate weights, all dissections were carried out in PBS and blotted on a Kimwipe immediately before weighing.

### Nuclei Dissociation of Brain Tissue

To test for energetic trade-offs mediating brain evolution through shifts in cellular composition we determined the number of neurons and glial cells using an adaptation of the isotropic fractionator method (Herculano-Houzel & Lent, 2005). This approach provides a rapid and accurate count of neuronal and glial cells in whole brains by dissociating nuclei and staining with a neuron-specific marker (Bahney & von Bartheld, 2014; Miller et al., 2014; Ngwenya et al., 2017). Briefly, whole brains were fixed in 1% formaldehyde for 10 min. Unused formaldehyde was removed by adding Glycine (0.125 M) and incubating for 5 minutes, at which point they were centrifuged (5 min at 1100 g, 4°C) and liquid was removed. To extract nuclei from the fixed cells the tissue was incubated on ice for 30 min in 1 mL of NF1 (10 mM Tris-HCL, 1 mM EDTA, 5 mM MgCl_2_, 0.1 M sucrose, 0.5% Triton X-100, diH_2_O), and homogenized using a Dounce tissue grinder (Kimble, Vineland, NJ, USA) using both the loose and tight pestles. To remove detritus the nuclei were passed through a 40 μm mini strainer (pluriSelect, El Cajon, CA, USA), centrifuged (5 min at 1600g, 4°C) and resuspended in 200 μL NF1. To visualize nuclei, they were stained by adding DAPI (4,6-Diamidino-2-Phenylindole, Dihydrochloride) (10% of the total volume) and incubated for 10 min at 4°C. Nuclei were then pelleted (5 min at 1600g, 4°C) and resuspended in 300μL NF1 before dividing into aliquots to be used for quantifying either the total number of cells in the brain (100 μL aliquot) or neuron/glia ratio (200μL aliquot).

### Quantification of Total Brain Cells

To quantify the total number of cells in the brain, we homogenized each sample by vortexing vigorously and imaging ten aliquots (10 μL each) using a Cell Countess 3 FL with DAPI 2.0 EVOS light cube (Invitrogen, Waltham, MA, USA). Quantification of each aliquot was performed by manually counting nondamaged nuclei using the multi-point feature in FIJI. Outlier aliquot were identified and discarded following best practice for Isotropic fractionation (Herculano-Houzel & Lent, 2005; Marhounová et al., 2019). In brief, we determined the coefficient of variation across aliquots for each individual, and if the coefficient of variation was ≥ 0.10 we considered the aliquot furthest from the mean to be an outlier and removed it (for the nosiest sample, only three of the ten aliquots had to be discarded). The total number of cells in the brain was then calculated by multiplying the mean (across aliquots) by the dilution factor.

### Quantifying Ratio of Neurons to Glial Cells

To quantify the ratio of neurons to glial cells, we used a monoclonal antibody (NeuN) to selectively label neuronal nuclei (Mullen et al., 1992). The 200μL suspension of dissociated brain nuclei was washed and resuspended in PBS then incubated overnight with primary antibody (1: 300 Rabbit antiNeuN antibody; abcam, Cambridge, United Kingdom). The next day, nuclei were washed in PBS, resuspended (90% PBS, 10% Bovine Serum Albumin), and incubated for two hours with DAPI (10 ug/mL) and secondary antibody (1: 400 Alexa Fluor® 488 AffiniPure Donkey Anti-Rabbit IgG; Jackson ImmunoResearch, West Grove, PA, USA). After incubation, nuclei were pelleted and resuspended in 100 μL of PBS.

We measured neuron to glia ratio in six aliquots (10 μL each) for each sample by first imaging all nuclei (both neuronal and glial) then imaging only neuronal nuclei. To do this we imaged aliquots on a Cell Countess 3 FL first under a DAPI 2.0 EVOS light cube to identify all nuclei, then under a GFP 2.0 EVOS light cube to identify only nuclei with signal from NeuN antibody (Invitrogen, Waltham, MA, USA). Nuclei were classified as neuronal if they stained with both DAPI and Alexa Fluor 488 and glial if they only stained with DAPI. The neuron/glia ratio was determined for each aliquot by dividing the number of neuronal nuclei by the total number of nuclei (neuronal + glial).

### Brain and Gonad Investment Differences Across Reproductive Tactics

To account for potential differences in absolute tissue size that are just due to allometric scaling, we corrected absolute tissue size (brain weight and gonad weight) by body size (standard length) as the residual of tissue on body length following Sukhum et al (2016):

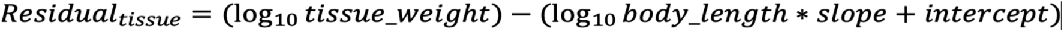

We then ran all analyses for each tissue on both raw weights and residual weights. Initial three-way ANOVAs (model: trait ∼ sex*species + species: morph) revealed significant effects of sex in every trait (Supplementary Tables S5, S9), therefore we conducted analyses independently for each sex.

To identify whether tissues differ between males utilizing different reproductive tactics, we compared male gonad weight, brain weight and neuron/glia ratio across species and morphs using two-way ANOVAs using the model (trait ∼ species + species: morph). We found strong differences for each of the male traits, so we ran Tukey’s HSD to make pairwise comparisons (R package: emmeans (Lenth, 2024)).

Despite the strong differences in brain size between males, a one-way ANOVA of female brain size revealed no differences across species (Supplementary Tables S8, S12).

### Evolutionary Trade-offs between Brain and Gonad

To test whether the relationship between brain traits and male gonad size differs across species, we grouped all morphs of *P. parae* and compared species using a general linear model (LM) in which the dependent variable was brain trait (either residual brain weight or neuron/glia ratio) and the independent variable was residual testes weight for each species (model: brain_trait ∼ gonad_size*species) (R package: lme4; Bates et al., 2015). To compare the slope and intercept we used estimated marginal means (EMMs), adjusting for multiple comparisons with Tukey’s method (R package: emmeans; Lenth, 2024).

Next, we focused on the relationship between brain traits and male gonad size in the individual male morphs of *P. parae*, by testing if morphs differ from one another and/or the closely related species (model: brain_trait ∼ gonad_size*male_type). Slope and intercept were again compared with EMM.

All statistical analyses were performed in the R statistical environment (R Core Team, 2023) and details can be found in the Supplemental Material. R code was generated with the help of ChatGPT (OpenAI, 2023) and graphs were produced in GraphPad Prism (*GraphPad Prism*, 2025).

## Results

### Gonad Size Differs Between Male Reproductive Tactics

We found gonad size differs by morph and species both when comparing raw weight (Table S1; ANOVA: df=2, F_2,45_=9.59, P <0.001) and when correcting for body size (Table S3; ANOVA: df=2, F_2,45_=15.65, P <0.001). Comparing across the male morphs of *P. parae* we found the immaculata morph has larger testes than both the melanzona and parae morphs, remarkably this relationship held both when correcting for body size (Fig. 1B, Table S4; Tukey: melanzona: t ratio= 3.06, p=0.029; parae: t ratio= 5.59, p<0.001), and even when comparing raw testes sizes, the immaculata gonads were 1.43 times larger than the parae morph and 1.30 times larger than the melanzona morph (Fig. 1B, Table S2; melanzona: t ratio= 3.27, p=0.017; parae: t ratio=4.16, p=0.001). The immaculata morph also had larger testes than both of the sister taxa, which is consistent with selection for increased gonad size in the immaculata morph (Fig. 1B, Table S4; *P. picta*: t ratio= 3.24, p=0.018; *P. reticulata*: t ratio=4.88 p<0.001).

### Brain Size Differs Between Male Reproductive Tactics

Brain size also differed between species and morphs both when comparing raw weight (Table S6; ANOVA: df=2, F_2,45_=8.96, P <0.001) and when correcting for body size (Table S10; ANOVA: df=2, F_2,45_=10.67, P <0.001). The brain size of the immaculata morph was 0.77 times smaller than the other morphs both when comparing raw weight (Fig. 1C, Table S7; melanzona: t ratio=-2.99, p=0.034; parae: t ratio=-4.09, p=0.002), and after correcting for their body size (Fig. 1C, Table S11; melanzona: t ratio= -4.08, p=0.002; parae: t ratio= -3.91, p=0.003). The relative brain size of the immaculata morph was also smaller than males of both sister taxa (Fig. 1C, Table S11; *P. picta*: t ratio= -7.47, p<0.001; *P. reticulata*: t ratio= -3.99, p<0.002).

Males of *P. picta* had the largest brains both in raw weight (Fig. 1C, Table S7; *P. parae* morphs: parae: t ratio= -3.46, p=0.01; melanzona: t ratio= -4.55, p<0.001, immaculata: t ratio= -7.55, p=0.012; *P. reticulata*: t ratio= 7.84, p<0.001) and when correcting for body size (Fig. 1C, Table S11; *P. parae* morphs: parae: t ratio= -3.55, p=0.008; melanzona: t ratio= -3.39, p<0.001, immaculata: p=0.012; *P. reticulata*: t ratio= 3.47, p=0.01).

### Neuron/Glia Ratio Does Not Differ Between Male Reproductive Tactics

The ratio of neurons to glial cells in the brain did not differ between male morphs of *P. parae* (Table S14; ANOVA: F_2,45_=2.81, P =0.071). However, neuron/glial ratio did differ between species (Table S14; ANOVA: df=2, F_2,45_=10.52, P <0.001), with both *P. picta* and *P. reticulata* males having a higher neuron/glia ratio compared to *P. parae* males (Fig. 1D, Table S15; *P. picta*: p<0.001, *P. reticulata*: p=0.018).

### Tissue Trade-offs Differ Between Male Reproductive Tactics

Consistent with an energetic trade-off, there was a negative correlation between brain and gonad sizes in both of the species utilizing flexible mating strategies, *P. picta* (Table S18; slope= -0.44, p = 0.027) and *P. reticulata* (slope= -0.60, p = 0.037) (Fig. 2A, Table S18; LM: F_5,44_= 8.17, P< 0.001). However, when combining the specialized morphs of *P. parae*, there was no relationship between these traits (Fig. 2A; slope= -0.12, p=0.18), though this slope was not different than either species (*P. picta*: p= 0.29, *P. reticulata*: p=0.24).

**Figure 2.**
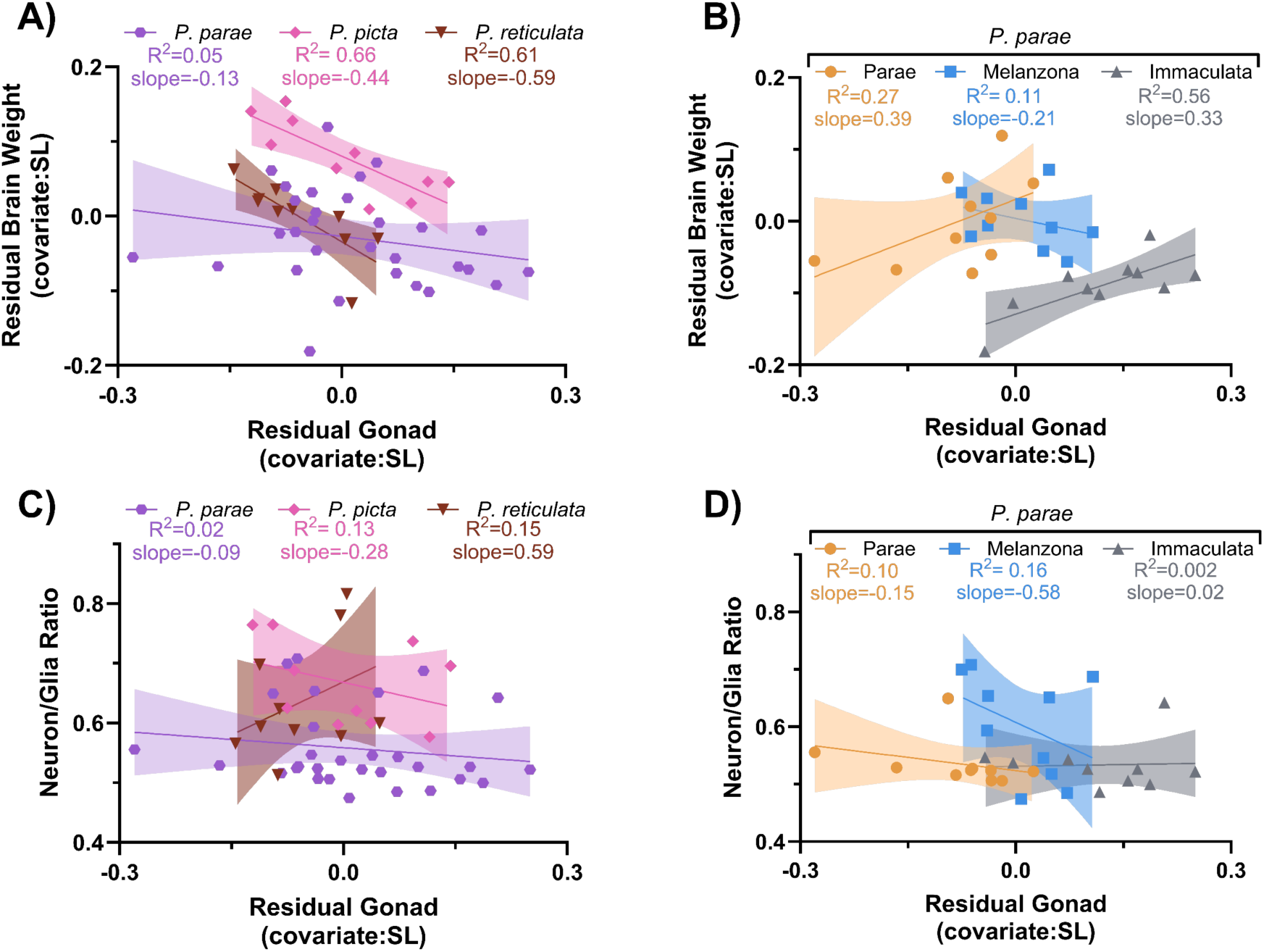
The negative relationship between male brain size and gonad size observed in all three species is consistent with a trade-off (A). However, while there was a strong correlation in both *P. picta* and *P. reticulata* (R^2^= 0.66 and 0.61) the relationship was weak in *P. parae* (R^2^=0.05). Yet when examining the male morphs of *P. parae* independently (B), we see the negative correlation was driven by morph differences, primarily from the large gonads and small brains of the immaculata morph. Despite the potential for energetic trade-offs to be mediated through shifts in neuron/glia ratios, we found no correlation with gonad size in any of the three species (C) or morphs (D). General linear models were used followed by Tukey comparison with the shaded regions indicating the 95% confidence interval. R^2^ indicates the fit to the regression.

Surprisingly, when examining the morphs of *P. parae* independently we found relationships that were inconsistent with brain and gonad sizes being mediated through an energetic trade-off. Rather than a negative relationship, we found gonad size was positively correlated with brain size in both the immaculata (Fig. 2B, Table S19; LM: F_9,40_= 13.45, P< 0.001; slope= 0.33, p = 0.019) and parae morphs (slope= 0.38, p = 0.011). These positive relationships differed significantly from the sister species for both the parae morph (Table S19; with *P. picta*: p = 0.001, with *P. reticulata*: p = 0.023) and immaculata morph (with *P. picta*: p = 0.002, with *P. reticulata*: p = 0.033). Meanwhile, there was no relationship between brain size and gonad size in the melanzona morph (slope=-0.21, p=0.36).

We found no support for energetic trade-offs being mediated by changes to cell types as there was no relationship between residual gonad size and neuron/glia ratio across morphs or species (Fig. 2C&D, Table S20,S21; LM: F_9,40_= 3.32, P= 0.004).

### Female Brain Size Did Not Differ Across Species

Comparing brain size by sex and species revealed a significant effect of sex by species both on raw brain weight (Table S5; ANOVA: df=2, F_2,72_=6.14, P =0.003) and when correcting for body size (Table S9; ANOVA: df=2, F_2,72_=6.98, P =0.002). However, this affect was entirely due to variation in males since examining brain size of females across species revealed no difference in either raw brain weight (Fig. S2A, Table S8; ANOVA: df=2, F_2,27_=2.23, P =0.13) or when correcting by body size (Fig. S2B, Table S12; ANOVA: df=2, F_2,27_=0.04, P =0.96). The male-limited effects of species differences in brain size is consistent with male reproductive strategy having an effect on brain size. Meanwhile, the neuron/glia ratio of females also differed across species (Table S16; ANOVA: df=2, F_2,27_=5.72, P =0.008), where the neuron/glia ratio of *P. picta* was higher than both *P. parae* (Figure S2C, Table S17; t ratio=3.04, p=0.014) and *P. reticulata* (t ratio=2.81, p=0.024).

## Discussion

Male reproductive strategies often select for increased gonad size, which has been proposed to drive concomitant decreases in brain size through energetic trade-offs between tissues (Aiello & Wheeler, 1995; Heldstab et al., 2022; Lemaître et al., 2009). If such energetic trade-offs are what mediates the co-evolution of gonad and brain sizes, then in the absence of ecological differences larger gonads should always correspond with less energetically costly brains, both within and across species. We found males of the species *P. picta* and *P. reticulata* indeed showed a negative correlation between gonad and brain sizes. However, this negative relationship was not found in any of the male morphs of *P. parae*, and surprisingly, a positive relationship was observed in both the parae and immaculata morphs. This suggests that male reproductive tactics are not selecting on energetic trade-offs *per se,* but instead, there is correlated selection for large gonads and small brains. Below, we elaborate on what this lack of energetic trade-off means and how generalist versus specialist mating strategies could explain observed negative correlations between gonad size and brain size.

### No Energetic Trade-off Between Male Gonad and Brain

Across the three species, males exhibited an overall negative correlation between brain and gonad size (Fig. 2A). However, when examining the individual morphs of *P. parae* we found that increased gonad size was not always accompanied by concomitant decreases in brain size (Fig. 2B). The immaculata morph had both larger gonads and smaller brains compared to the other morphs. However, there was a positive relationship between brain and gonad size across individuals within the parae and immaculata morphs. This positive within-morph correlation is consistent with allometric scaling but is inconsistent with two traits being under an energetic trade-off.

We did recover a negative relationship between gonad size and brain size in both *P. picta* and *P. reticulata*. While it is possible this negative relationship between gonad and brain size is due to energetic trade-offs in these species, this seems less likely based on a selection experiment on brain size conducted in *P. reticulata*. After establishing selection lines of large brained and small brained *P. reticulata*, Kotrschal et al. (2015) found no difference in gonad size between lines. Interestingly, while this provides compelling support that a trade-off is not being driven through differences in energy allocation, it is still possible there is a trade-off being mediated through correlated selection since selection on one trait under a non-energetic trade-off will often result in a shift to the Y intercept but retain the slope (Roff & Fairbairn, 2007).

Brains are primarily energetically expensive due to the high metabolic demands of neuron cells, which are supported by glia cells (Attwell & Laughlin, 2001; Harris et al., 2012). We tested whether energetic trade-offs with gonad size were shaping brain evolution through shifts in the cellular composition but found no relationship between gonad size and neuron/glia ratio. Given that neuron/glia ratio appears to be a readily evolvable trait in other systems (Herculano-Houzel, 2014), the lack of a relationship between gonad size and neuron/glia ratio is consistent with the conclusion that gonads and brains are not directly competing for energy allocation.

While several studies have tested for energetic trade-offs between gonad size and brain evolution, only a couple have provided some support. These have been in swordtail fish, where only one of the two behavioural morphs showed support (Stec et al., 2023), and a comparative study across bat species (Pitnick et al., 2005). However, when reanalysed along with more comparisons across mammals, Lemaître et al. (2009) did not find a negatively relationship across bat species or other mammals, but rather only in echolocating bats. In fact, many studies have failed to find a negative relationship between brain and gonads (odontocetes; Kelley et al., 2014; frogs; Jin et al., 2015, Zhao et al., 2016, fish; Warren & Iglesias, 2012, mammals; Bordes et al., 2011; Husak et al., 2024; Lemaître et al., 2009; Schillaci, 2006). Our results, together with these previous studies, suggest that negative correlations between brain and gonad size are not due to differences in energetic allocation, but instead arise through highly specific constraints imposed by other forces such as differences in ecological niche (reviewed by Garland et al., 2022) or correlated selection (Agrawal, 2020; Grabowski et al., 2023).

### Corelated Selection for Large Gonads and Small Brains

When multiple traits contribute to fitness, selection will act to drive traits toward combinations that increase fitness through correlated selection (Lande, 1979; Turelli, 1985). Often alternative combinations of traits exist which can maximize fitness in different ways, resulting in multiple peaks on the fitness landscape (Whitlock & Phillips, 1995). When traits are not tightly linked, recombination can generate multiple trait combinations and the population will explore the fitness landscape (Wright, 1932). Conversely, when traits are genetically linked and recombination is limited, specific trait combinations can allow selection toward a single peak and the emergence of specialization (Futuyma & Moreno, 1988). Across the three species, males exhibited an overall negative correlation between gonad and brain sizes (Fig. 2A). However, this pattern was not observed within the morphs of *P. parae* (Fig. 2B), suggesting morphs have undergone selection for distinct, optimized trait combinations.

Unlike the specialized morphs of *P. parae*, both *P. picta* and *P. reticulata* exhibit more generalist reproductive strategies, where males flexibly employ both sneaking and display tactics rather than specializing in a single mating approach (Houde, 1997; Liley, 1966; Pilastro & Bisazza, 1999). Males vary across individuals in the degree to which they use each strategy (Liley, 1966; Reynolds et al., 1993), which suggests that the negative relationship we observed may be linked to this individual variability.

The male morphs of *P. parae* are inherited through alternative, non-recombining Y chromosome haplotypes (Lindholm et al., 2004, 2006), which provide large morph-specific regions of the genome upon which selection can act on each morph individually. This genetic architecture allows morphs to evolve toward alternative fitness peaks, reducing the overall trait correlation, as reflected by the lack of a negative correlation between brain and gonad size in *P. parae* (Fig. 2A). The immaculata morph, which relies on sneak copulations and sperm competition, is under selective pressure to increase gonad size (Dougherty et al., 2022; Hurtado-Gonzales & Uy, 2009). We found this obligate sneaker morph had larger gonads and smaller brains than either of the other morphs (Fig. 1), consistent with selection driving the immaculata morph toward a fitness peak favouring larger gonads, interestingly this seems to have been accompanied by correlated selection for smaller brains.

Multiple studies have documented negative correlations between brain size and other energetically costly organs, including gonads (bats: Pitnick et al., 2005; swordtail fish: Stec et al., 2023), muscle (birds: Isler & van Schaik, 2009), gut (Jin et al., 2015; Kotrschal et al., 2013), fat storage (cichlids: Tsuboi et al., 2016), and liver (toads: Yao et al., 2025). These negative relationships are often attributed to energetic trade-offs. However, as seen in *P. parae*, such correlations may not always be driven by direct constraints of energy allocation but could instead arise through correlated selection optimizing trait combinations for specific reproductive strategies. This highlights the need for models of brain size evolution to consider more than just energetic constraints when interpreting negative trait relationships across species.

## Conclusion

Our findings challenge the hypothesis that selection on mating strategy can drive brain size evolution through energetic trade-offs with gonad size. Instead, our results are consistent with negative correlations between gonad size and brain size being driven through correlated selection between traits. Specifically, while male gonad size negatively correlated with brain size in *P. picta* and *P. reticulata*, which utilize generalist reproductive strategies, the male morphs of *P. parae*, which specialize in alternative reproductive strategies, had a positive correlation. This suggests that male reproductive strategies may impose correlated selection that results in negative correlations between gonad and brain size. Understanding how trade-offs are realized within and between specialized strategies will provide deeper insights into the evolution of reproductive strategies and tissue allocation.

## Acknowledgements

We thank Drs. Anurag Agrawal, Kern Reeve, and Nora Prior for helpful feedback on the manuscript, along with all members of the Sandkam lab for discussions and assistance with fish care, especially Maximilliano Zuluaga Forero, Meridia Bryant, Dr. Ehren Bentz, Dr. Matthew Taves, Lynn Capani-Czebiniak and Sri Sai Akkineni. Finally, we thank our funding sources: Cornell University, NSF, and NIH. Research reported in this publication was supported by the National Institute Of General Medical Sciences of the National Institutes of Health under Award Number R35GM155164. The content is solely the responsibility of the authors and does not necessarily represent the official views of the National Institutes of Health.

