## Supplemental Materials for "Specialization of male reproductive tactics correspond with large gonads and small brains"

**Supplemental Figures and Tables for Male reproductive tactics drive correlated selection for large gonads with small brains**

Helen Stec[1], Grace Y. Zhang[1], Ben A. Sandkam[1]

Supplementary Figures

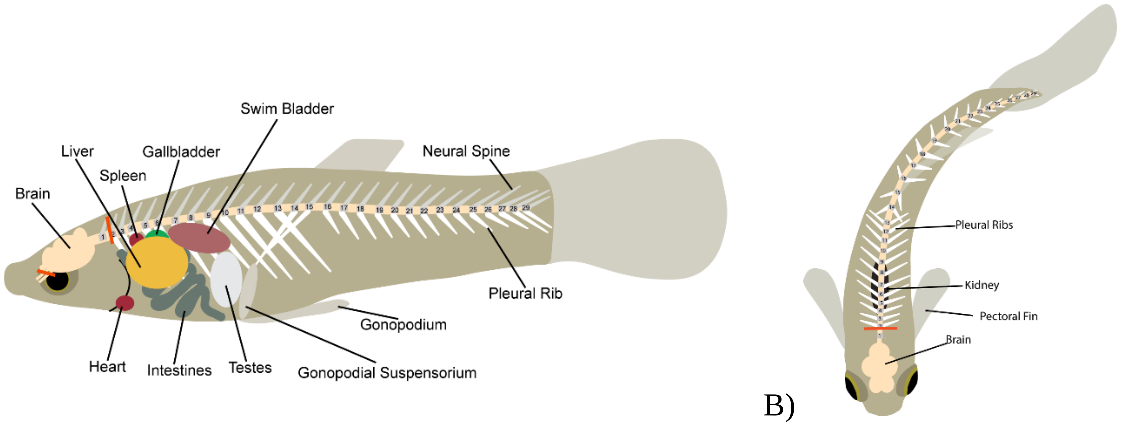


A) B)

**Figure S1**. Brain dissections were performed using precise landmarks to ensure weight differences were not due to inconsistent dissections. Diagrams of a male *P. reticulata* noting the location of the organs dissected and the landmarks used to dissect the brain with a (A) a sagittal view and (B) a dorsal view. The red lines indicate the landmarks used to dissect out the brain. Brain stems were cut anterior to the first pleural rib, found on the second vertebrae. The optic nerves were cut at their connection point.

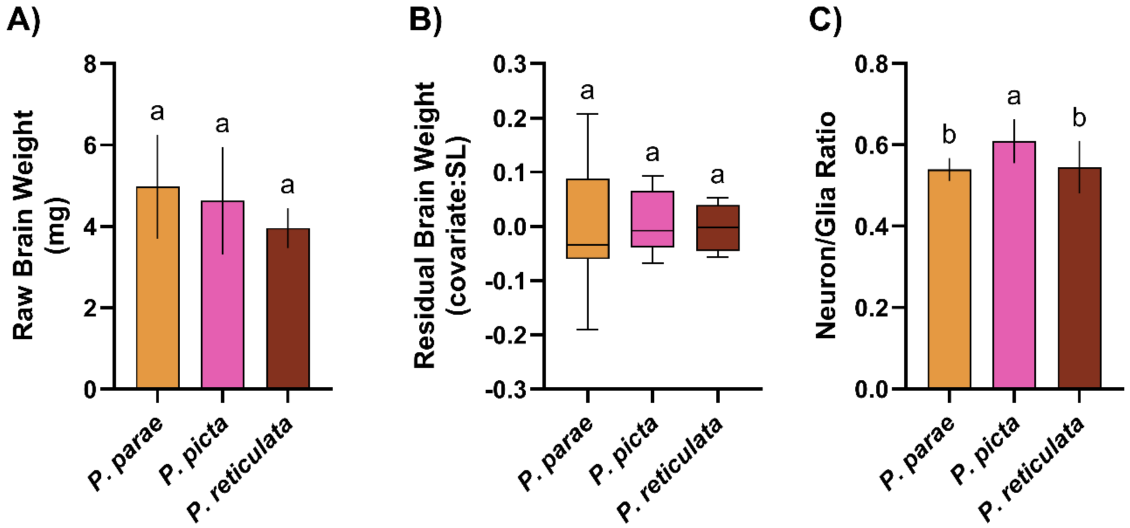


Fig. S2. **Female brain size** did not differ across *P. parae*, *P. picta*, and *P. reticulata*, both when comparing raw weights (A) or correcting for body size (B). Females of *P. picta* had higher neuron/glia ratios compared to females of both sister species (C). One-way ANOVAs followed by Tukey comparisons were used with error bars indicating SE, letters denote significant different groups ($p<0.05$).

*Supplementary Tables*

Table S1: **Male gonad weight (raw)** differs by species and morph within species as revealed by Two-way ANOVA. Significant p-values ($p<0.05$) in bold.

| Effect | df | Sum Sq | Mean Sq | F value | Pr (>F) |
|---|---|---|---|---|---|
| **Species** | 2 | 21.5 | 10.767 | 20.45 | **4.81e-07** |
| **Species:Morph** | 2 | 10.1 | 5.050 | 9.59 | **3.39e-04** |
| Residuals | 45 | 23.7 | 0.526 | NA | NA |


| Effect | df | Sum Sq | Mean Sq | F value | Pr (>F) |
|---|---|---|---|---|---|
| **Species** | 2 | 21.5 | 10.767 | 20.45 | **4.81e-07** |
| **Species:Morph** | 2 | 10.1 | 5.050 | 9.59 | **3.39e-04** |
| Residuals | 45 | 23.7 | 0.526 | NA | NA |


Table S2: Tukey Pairwise Comparisons reveal the **male gonad weight (raw)** of the Immaculata morph of *P. parae* differs from the other morphs and *P. reticulata*. Significant pairwise comparisons in bold and underlined ($p<0.05$).

| Contrast | Estimate | SE | df | t ratio | P value |
|---|---|---|---|---|---|
| **Immaculata [*P. parae*] – Melanzona [*P. parae*]** | **1.06** | **0.324** | **45** | **3.267** | **1.69e-02** |
| **Immaculata [*P. parae*] – Parae [*P. parae*]** | **1.35** | **0.324** | **45** | **4.160** | **1.27e-03** |
| Immaculata [*P. parae*] – *P. picta* | 0.57 | 0.324 | 45 | 1.757 | 4.11e-01 |
| **Immaculata [*P. parae*] – *P. reticulata*** | **2.37** | **0.324** | **45** | **7.304** | **3.56e-08** |
| Melanzona [*P. parae*] – Parae [*P. parae*] | 0.29 | 0.324 | 45 | 0.894 | 8.98e-01 |
| Melanzona [*P. parae*] – *P. picta* | -0.49 | 0.324 | 45 | -1.510 | 5.62e-01 |
| **Melanzona [*P. parae*] – *P. reticulata*** | **1.31** | **0.324** | **45** | **4.037** | **1.86e-03** |
| Parae [*P. parae*] – *P. picta* | -0.78 | 0.324 | 45 | -2.404 | 1.33e-01 |
| **Parae [*P. parae*] – *P. reticulata*** | **1.02** | **0.324** | **45** | **3.143** | **2.35e-02** |
| ***P. picta* – *P. reticulata*** | **1.80** | **0.324** | **45** | **5.547** | **1.40e-05** |

Table S3: **Male gonad weight (as residual of body size)** differs by species and morph within species as revealed by Two-way ANOVA. Significant p-values ($p<0.05$) in bold.

| Effect | df | Sum Sq | Mean Sq | F value | Pr (>F) |
|---|---|---|---|---|---|
| Species | 2 | 0.0393 | 0.01964 | 3.0 | 5.98e-02 |
| **Species: Morph** | 2 | 0.2048 | 0.10239 | 15.6 | **6.94e-06** |
| Residuals | 45 | 0.2945 | 0.00654 | NA | NA |

Table S4: Tukey Pairwise Comparisons reveal **male gonad weight (as residual of body size)** of the Immaculata morph of *P. parae* differs from the other morphs and *P. reticulata.* Significant pairwise comparisons in bold and underlined ($p<0.05$).

| Contrast | Estimate | SE | df | t ratio | P value |
|---|---|---|---|---|---|
| **Immaculata [*P. parae*] – Melanzona [*P. parae*]** | **0.11078** | **0.0362** | **45** | **3.062** | **2.89e-02** |
| **Immaculata [*P. parae*] – Parae [*P. parae*]** | **0.20206** | **0.0362** | **45** | **5.585** | **1.23e-05** |
| **Immaculata [*P. parae*] – *P. picta*** | **0.11704** | **0.0362** | **45** | **3.235** | **1.84e-02** |
| **Immaculata [*P. parae*] – *P. reticulata*** | **0.17645** | **0.0362** | **45** | **4.877** | **1.30e-04** |
| Melanzona [*P. parae*] – Parae [*P. parae*] | 0.09128 | 0.0362 | 45 | 2.523 | 1.03e-01 |
| Melanzona [*P. parae*] – *P. picta* | 0.00626 | 0.0362 | 45 | 0.173 | 1.00e+00 |
| Melanzona [*P. parae*] – *P. reticulata* | 0.06567 | 0.0362 | 45 | 1.815 | 3.78e-01 |
| Parae [*P. parae*] – *P. picta* | -0.08502 | 0.0362 | 45 | -2.350 | 1.48e-01 |
| Parae [*P. parae*] – *P. reticulata* | -0.02561 | 0.0362 | 45 | -0.708 | 9.54e-01 |
| *P. picta* – *P. reticulata* | 0.05941 | 0.0362 | 45 | 1.642 | 4.79e-01 |

Table S5: **Brain weight (raw)** differs by sex, species, and morph within species as revealed by a three-way ANOVA. Significant terms in bold (p<0.05) .

| Effect | df | Sum Sq | Mean Sq | F value | Pr (>F) |
|---|---|---|---|---|---|
| **Sex** | 1 | 10.93 | 10.925 | 17.43 | **8.25e-05** |
| **Species** | 2 | 16.05 | 8.026 | 12.80 | **1.75e-05** |
| **Sex: Species** | 2 | 7.70 | 3.849 | 6.14 | **3.45e-03** |
| **Species: Morph** | 2 | 5.04 | 2.521 | 4.02 | **2.21e-02** |
| Residuals | 72 | 45.14 | 0.627 | NA | NA |

Table S6: **Male brain weight (raw)** differs by species and by morphs of *P. parae* as revealed by two-way ANOVA. Significant terms in bold (p<0.05).

| Effect | df | Sum Sq | Mean Sq | F value | Pr (>F) |
|---|---|---|---|---|---|
| **Species** | 2 | 18.38 | 9.189 | 32.66 | **1.73e-09** |
| **Species: Morph** | 2 | 5.04 | 2.521 | 8.96 | **5.30e-04** |
| Residuals | 45 | 12.66 | 0.281 | NA | NA |

Table S7: Tukey Pairwise Comparisons show the **brain weight (raw)** of the immaculata morph of *P. parae* differs from the other two *P. parae* morphs and *P. reticulata*. Significant pairwise comparisons in bold and underlined (p<0.05).

| Contrast | Estimate | SE | df | t ratio | P value |
|---|---|---|---|---|---|
| **Immaculata [*P. parae*] – Melanzona [*P. parae]*** | **-0.71** | **0.237** | **45** | **-2.993** | **3.44e-02** |
| **Immaculata [*P. parae*] – Parae [*P. parae*]** | **-0.97** | **0.237** | **45** | **-4.089** | **1.58e-03** |
| **Immaculata [*P. parae*] – *P. picta*** | **-1.79** | **0.237** | **45** | **-7.546** | **1.57e-08** |
| Immaculata [*P. parae*] – *P. reticulata* | 0.07 | 0.237 | 45 | 0.295 | 9.98e-01 |
| Melanzona [*P. parae*] – Parae [*P. parae*] | -0.26 | 0.237 | 45 | -1.096 | 8.08e-01 |
| **Melanzona [*P. parae*] – *P. picta*** | **-1.08** | **0.237** | **45** | **-4.553** | **3.71e-04** |
| **Melanzona [*P. parae*] – *P. reticulata*** | **0.78** | **0.237** | **45** | **3.288** | **1.60e-02** |
| **Parae [*P. parae*] – *P. picta*** | **-0.82** | **0.237** | **45** | **-3.457** | **1.01e-02** |
| **Parae [*P. parae*] – *P. reticulata*** | **1.04** | **0.237** | **45** | **4.384** | **6.34e-04** |
| ***P. picta* – *P. reticulata*** | **1.86** | **0.237** | **45** | **7.841** | **5.80e-09** |

Table S8: **Female brain weight (raw)** does not differ by species as revealed by one-way ANOVA.

| Effect | df | Sum Sq | Mean Sq | F value | Pr (>F) |
|---|---|---|---|---|---|
| Species | 2 | 5.37 | 2.69 | 2.23 | 0.127 |
| Residuals | 27 | 32.48 | 1.20 | NA | NA |

Table S9: Three-way ANOVA shows **brain weight (as residual of body size)** differs by species, sex within species, and morph within species. Significant effects in bold (p<0.05).

| Effect | df | Sum Sq | Mean Sq | F value | Pr (>F) |
|---|---|---|---|---|---|
| Sex | 1 | 1.32e-06 | 1.32e-06 | 0.000347 | 0.98519 |
| **Species** | 2 | 5.77e-02 | 2.89e-02 | 7.564642 | **0.00104** |
| **Sex: Species** | 2 | 2.91e-02 | 1.45e-02 | 3.811878 | **0.02670** |
| **Species: Morph** | 2 | 5.33e-02 | 2.66e-02 | 6.978885 | **0.00170** |
| Residuals | 72 | 2.75e-01 | 3.82e-03 | NA | NA |

Table S10: Two-way ANOVA reveals the **brain weight of males (as residual of body size)** differs by species and by morphs of *P. parae*. Significant effects in bold (p<0.05).

| Effect | df | Sum Sq | Mean Sq | F value | Pr (>F) |
|---|---|---|---|---|---|
| **Species** | 2 | 0.0863 | 0.0432 | 17.3 | **2.68e-06** |
| **Species: Morph** | 2 | 0.0533 | 0.0266 | 10.7 | **1.61e-04** |
| Residuals | 45 | 0.1123 | 0.0025 | NA | NA |

Table S11: Tukey Pairwise Comparisons show the **male brain weight (as residual of body size)** of the immaculata morph of *P. parae* differs from the other two *P. parae* morphs and males of both sister species. Significant pairwise comparisons in bold and underlined (p<0.05).

| Contrast | Estimate | SE | df | t ratio | P value |
|---|---|---|---|---|---|
| **Immaculata [*P. parae*] – Melanzona [*P. parae*]** | **-0.09107** | **0.0223** | **45** | **-4.0763** | **1.65e-03** |
| **Immaculata [*P. parae*] – Parae [*P. parae*]** | **-0.08762** | **0.0223** | **45** | **-3.9221** | **2.63e-03** |
| **Immaculata [*P. parae*] – *P. picta*** | **-0.16681** | **0.0223** | **45** | **-7.4667** | **2.05e-08** |
| **Immaculata [*P. parae*] – *P. reticulata*** | **-0.08919** | **0.0223** | **45** | **-3.9925** | **2.12e-03** |
| Melanzona [*P. parae*] – Parae [*P. parae*] | 0.00344 | 0.0223 | 45 | 0.1542 | 1.00e+00 |
| **Melanzona [*P. parae*] – *P. picta*** | **-0.07575** | **0.0223** | **45** | **-3.3905** | **1.21e-02** |
| Melanzona [*P. parae*] – *P. reticulata* | 0.00187 | 0.0223 | 45 | 0.0838 | 1.00e+0 |
| **Parae [*P. parae*] – *P. picta*** | **-0.07919** | **0.0223** | **45** | **-3.5446** | **7.88e-03** |
| Parae [*P. parae*] – *P. reticulata* | -0.00157 | 0.0223 | 45 | -0.0704 | 1.00e+00 |
| ***P. picta* – *P. reticulata*** | **0.07762** | **0.0223** | **45** | **3.4743** | **9.60e-03** |

Table 12: **Female brain weight (as residual of body size)** does not differ by species as revealed by one-way ANOVA.

| Effect | df | Sum Sq | Mean Sq | F value | Pr (>F) |
|---|---|---|---|---|---|
| Species | 2 | 0.000515 | 0.000257 | 0.0428 | 0.958 |
| Residuals | 27 | 0.162518 | 0.006019 | NA | NA |

Table S13: Three-way ANOVA revealed the **Neuron/Glia ratio** differs by sex, species, and morph within species. Significant effects in bold ($p<0.05$).

| Effect | df | Sum Sq | Mean Sq | F value | Pr (>F) |
|---|---|---|---|---|---|
| **Sex** | 1 | 0.0172 | 0.01718 | 4.01 | **4.91e-02** |
| **Species** | 2 | 0.1247 | 0.06233 | 14.53 | **4.99e-06** |
| Sex: Species | 2 | 0.0166 | 0.00831 | 1.94 | 1.52e-01 |
| **Species: Morph** | 2 | 0.0298 | 0.01488 | 3.47 | **3.65e-02** |
| Residuals | 72 | 0.3088 | 0.00429 | NA | NA |

Table S14: Two-way ANOVA revealed the **Neuron/Glia ratio of males** differs by species, but not by morphs of *P. parae*. Significant effects in bold ($p<0.05$).

| Effect | df | Sum Sq | Mean Sq | F value | Pr (>F) |
|---|---|---|---|---|---|
| **Species** | 2 | 0.1114 | 0.0557 | 10.52 | **0.000179** |
| Species: Morph | 2 | 0.0298 | 0.0149 | 2.81 | 0.070897 |
| Residuals | 45 | 0.2384 | 0.0053 | NA | NA |

Table S15: Tukey Pairwise Comparison shows **Neuron/Glia ratio** of males of *P. parae* have a lower ratio compared to the sister species *P. picta* and *P. reticulata*. Since morph did not have an effect in the two-way ANOVA (Table S14), morph to species pairwise comparisons should be considered with caution. Significant pairwise comparisons in bold and underlined ($p<0.05$).

| Contrast | Estimate | SE | df | t ratio | P value |
|---|---|---|---|---|---|
| ***P. parae* – *P. picta*** | **-0.1099** | **0.0276** | **47** | **-3.984** | **0.000674** |
| ***P. parae* – *P. reticulata*** | **-0.0785** | **0.0276** | **47** | **-2.847** | **0.017584** |
| *P. picta* – *P. reticulata* | 0.0314 | 0.0338 | 47 | 0.928 | 0.625238 |
| Immaculata [*P. parae*] – Melanzona [*P. parae]* | -0.06796 | 0.0326 | 45 | -2.0879 | 0.24308 |
| Immaculata [*P. parae*] – Parae [*P. parae*] | -0.00237 | 0.0326 | 45 | -0.0728 | 0.99999 |
| **Immaculata [*P. parae*] – *P. picta*** | **-0.13334** | **0.0326** | **45** | **-4.0964** | **0.00155** |
| **Immaculata [*P. parae*] – *P. reticulata*** | **-0.10198** | **0.0326** | **45** | **-3.1329** | **0.02411** |
| Melanzona [*P. parae*] – Parae [*P. parae*] | 0.06559 | 0.0326 | 45 | 2.0151 | 0.27554 |
| Melanzona [*P. parae*] – *P. picta* | -0.06538 | 0.0326 | 45 | -2.0085 | 0.27863 |
| Melanzona [*P. parae*] – *P. reticulata* | -0.03402 | 0.0326 | 45 | -1.0450 | 0.83302 |
| **Parae [*P. parae*] – *P. picta*** | **-0.13097** | **0.0326** | **45** | **-4.0236** | **0.00193** |
| **Parae [*P. parae*] – *P. reticulata*** | **-0.09961** | **0.0326** | **45** | **-3.0601** | **0.02908** |
| *P. picta* – *P. reticulata* | 0.03136 | 0.0326 | 45 | 0.9635 | 0.86994 |

Table S16: One-way ANOVA revealed the **Neuron/Glia ratio of females** differs by species. Significant pairwise comparisons in bold (p<0.05).

| Effect | df | Sum Sq | Mean Sq | F value | Pr (>F) |
|---|---|---|---|---|---|
| **Species** | 2 | 0.0298 | 0.01492 | 5.72 | **0.00849** |
| Residuals | 27 | 0.0704 | 0.00261 | NA | NA |

Table S17: Tukey Pairwise Comparison shows **Neuron/Glia ratio** of females differs with *P. picta* females having a higher ratio compared to sister species *P. parae* and *P. reticulata*. Significant pairwise comparisons in bold and underlined (p<0.05).

| Contrast | Estimate | SE | df | t ratio | P value |
|---|---|---|---|---|---|
| ***P. parae – P. picta*** | **-0.06932** | **0.0228** | **27** | **-3.035** | **0.0141** |
| *P. parae – P. reticulata* | -0.00514 | 0.0228 | 27 | -0.225 | 0.9725 |
| ***P. picta – P. reticulata*** | **0.06419** | **0.0228** | **27** | **2.810** | **0.0239** |

Table S18: General linear model comparing **across species** the **brain weight of males (as residual of body size)** by **gonad weight of males (as residual of body size)** reveals a negative correlation in *P. picta* and *P. reticulata*. When combing the morphs of *P. parae* together, we see no correlation. Below is the linear model and follow up comparison of slope (compared to 0 and pairwise comparisons) and intercept (pairwise comparisons). Significant effects in bold and underlined (p<0.05).

| Model Results (residual brain weight ~ residual gonad weight*Species) | | | | |
|---|---|---|---|---|
| Effect | Estimate | Std. Error | t value | P value |
| **Linear Model** | | | | |
| **(Intercept)** | **-0.02543** | **0.0101** | **-2.528** | **1.51e-02** |
| Residual Gonad Weight (ResGL) | -0.12028 | 0.0880 | -1.366 | 1.79e-01 |
| ***P. picta*** | **0.10710** | **0.0200** | **5.363** | **2.89e-06** |
| *P. reticulata* | -0.00551 | 0.0252 | -0.219 | 8.28e-01 |
| ResGL: *P. picta* | -0.32710 | 0.2142 | -1.527 | 1.34e-01 |
| ResGL: *P. reticulata* | -0.47959 | 0.2920 | -1.643 | 1.08e-01 |
| **Slope to 0** | | | | |
| *P. parae* | -0.120 | 0.088 | -1.37 | 0.1789 |
| ***P. picta*** | **-0.447** | **0.195** | **-2.29** | **0.0268** |
| ***P. reticulata*** | **-0.600** | **0.278** | **-2.15** | **0.0367** |
| **Pairwise Slope Comparisons** | | | | |
| *P. parae* – *P. picta* | 0.327 | 0.214 | 1.527 | 0.288 |
| *P. parae* – *P. reticulata* | 0.480 | 0.292 | 1.643 | 0.239 |
| *P. picta* – *P. reticulata* | 0.152 | 0.340 | 0.448 | 0.895 |
| **Pairwise Intercept Comparisons** | | | | |
| ***P. parae* – *P. picta*** | **-0.10710** | **0.0200** | **-5.363** | **8.52e-06** |
| *P. parae* – *P. reticulata* | 0.00551 | 0.0252 | 0.219 | 9.74e-01 |
| ***P. picta* – *P. reticulata*** | **0.11261** | **0.0288** | **3.908** | **9.11e-04** |

Table S19: General linear model comparing **across male type** the **brain weight of males (as residual of body size)** by **gonad weight of males (as residual of body size)** reveals the parae and immaculata morphs of *P. parae* have a positive correlation. Below is the linear model and follow up comparison of slope (compared to 0 and pairwise comparisons) and intercept (pairwise comparisons). Significant effects in bold and underlined ($p<0.05$).

| Model Results (residual brain weight ~ residual gonad weight*male type) | | | | |
|---|---|---|---|---|
| Effect | Estimate | Std. Error | t value | P value |
| **Linear Model** | | | | |
| **(Intercept)** | **-0.1286** | **0.0213** | **-6.022** | **4.41e-07** |
| **Residual Gonad Weight (ResGL)** | **0.3424** | **0.1427** | **2.399** | **2.12e-02** |
| **Melanzona [*P. parae]*** | **0.1346** | **0.0248** | **5.420** | **3.08e-06** |
| **Parae [*P. parae*]** | **0.1603** | **0.0276** | **5.800** | **9.02e-07** |
| ***P. picta*** | **0.2102** | **0.0247** | **8.494** | **1.71e-10** |
| ***P. reticulata*** | **0.0976** | **0.0271** | **3.597** | **8.76e-04** |
| **ResGL: Melanzona [*P. parae]*** | **-0.5393** | **0.2571** | **-2.098** | **4.23e-02** |
| ResGL: Parae [*P. parae*] | 0.0437 | 0.2089 | 0.209 | 8.35e-01 |
| **ResGL: *P. picta*** | **-0.7897** | **0.2011** | **-3.927** | **3.31e-04** |
| **ResGL: *P. reticulata*** | **-0.9422** | **0.2473** | **-3.809** | **4.70e-04** |
| **Slope to 0** | | | | |
| **Immaculata [*P. parae*]** | **0.342** | **0.143** | **2.399** | **0.02117** |
| Melanzona [*P. parae*] | -0.197 | 0.214 | -0.921 | 0.36269 |
| **Parae [*P. parae*]** | **0.386** | **0.153** | **2.531** | **0.01539** |
| ***P. picta*** | **-0.447** | **0.142** | **-3.158** | **0.00302** |
| ***P. reticulata*** | **-0.600** | **0.202** | **-2.969** | **0.00503** |
| **Pairwise Slope Comparisons** | | | | |
| Immaculata [*P. parae*] – Melanzona [*P. parae]* | 0.5393 | 0.257 | 2.098 | 0.24109 |
| Immaculata [*P. parae*] – Parae [*P. parae*] | -0.0437 | 0.209 | -0.209 | 0.99955 |
| **Immaculata [*P. parae*] – *P. picta*** | **0.7897** | **0.201** | **3.927** | **0.00289** |
| **Immaculata [*P. parae*] – *P. reticulata*** | **0.9422** | **0.247** | **3.809** | **0.00406** |
| Melanzona [*P. parae*] – Parae [*P. parae*] | -0.5830 | 0.263 | -2.220 | 0.19349 |
| Melanzona [*P. parae*] – *P. picta* | 0.2505 | 0.257 | 0.976 | 0.86422 |
| Melanzona [*P. parae*] – *P. reticulata* | 0.4030 | 0.294 | 1.370 | 0.65007 |
| **Parae [*P. parae*] – *P. picta*** | **0.8335** | **0.208** | **4.004** | **0.00232** |
| **Parae [*P. parae*] – *P. reticulata*** | **0.9860** | **0.253** | **3.895** | **0.00318** |
| *P. picta* – *P. reticulata* | 0.1525 | 0.247 | 0.618 | 0.97137 |
| **Pairwise Intercept Comparisons** | | | | |
| **Immaculata [*P. parae*] – Melanzona [*P. parae]*** | **-0.1346** | **0.0248** | **-5.42** | **2.93e-05** |
| **Immaculata [*P. parae*] – Parae [*P. parae*]** | **-0.1603** | **0.0276** | **-5.80** | **8.66e-06** |
| **Immaculata [*P. parae*] – *P. picta*** | **-0.2102** | **0.0247** | **-8.49** | **1.69e-09** |
| **Immaculata [*P. parae*] – *P. reticulata*** | **-0.0976** | **0.0271** | **-3.60** | **7.39e-03** |
| Melanzona [*P. parae*] – Parae [*P. parae*] | -0.0257 | 0.0217 | -1.19 | 7.60e-01 |
| **Melanzona [*P. parae*] – *P. picta*** | **-0.0756** | **0.0178** | **-4.24** | **1.16e-03** |
| Melanzona [*P. parae*] – *P. reticulata* | 0.0370 | 0.0210 | 1.76 | 4.10e-01 |
| Parae [*P. parae*] – *P. picta* | -0.0499 | 0.0216 | -2.31 | 1.62e-01 |
| Parae [*P. parae*] – *P. reticulata* | 0.0627 | 0.0243 | 2.58 | 9.26e-02 |
| ***P. picta* – *P. reticulata*** | **0.1126** | **0.0209** | **5.38** | **3.29e-05** |

Table 20: General linear model comparing **across species** the **neuron/glia ratio of males (as residual of body size) by gonad weight of males (as residual of body size)** reveals no correlation in *P. parae* (morphs combined), *P. picta* and *P. reticulata*. Below is the linear model and follow up comparison of slope (compared to 0 and pairwise comparisons) and intercept (pairwise comparisons). Significant effects in bold and underlined (p<0.05).

| Model Results (Neuron/Glia ratio ~ residual gonad weight*Species) | | | | |
|---|---|---|---|---|
| Effect | Estimate | Std. Error | t value | P value |
| **Linear Model** | | | | |
| (Intercept) | 0.5587 | 0.0139 | 40.326 | 2.18e-36 |
| Residual Gonad Weight (ResGL) | -0.0927 | 0.1212 | -0.765 | 4.49e-01 |
| *P. picta* | 0.1095 | 0.0275 | 3.982 | 2.53e-04 |
| *P. reticulata* | 0.1064 | 0.0347 | 3.069 | 3.67e-03 |
| ResGL: *P. picta* | -0.1889 | 0.2949 | -0.640 | 5.25e-01 |
| ResGL: *P. reticulata* | 0.6268 | 0.4021 | 1.559 | 1.26e-01 |
| **Slope to 0** | | | | |
| *P. parae* | -0.0927 | 0.121 | -0.765 | 0.449 |
| *P. picta* | -0.2816 | 0.269 | -1.047 | 0.301 |
| *P. reticulata* | 0.5341 | 0.383 | 1.393 | 0.171 |
| **Pairwise Slope Comparisons** | | | | |
| *P. parae* – *P. picta* | 0.189 | 0.295 | 0.64 | 0.799 |
| *P. parae* – *P. reticulata* | -0.627 | 0.402 | -1.56 | 0.274 |
| *P. picta* – *P. reticulata* | -0.816 | 0.468 | -1.74 | 0.201 |
| **Pairwise Intercept Comparisons** | | | | |
| ***P. parae* – *P. picta*** | **-0.10951** | **0.0275** | **-3.9821** | **0.000726** |
| ***P. parae* – *P. reticulata*** | **-0.10643** | **0.0347** | **-3.0692** | **0.010060** |
| *P. picta* – *P. reticulata* | 0.00308 | 0.0397 | 0.0776 | 0.996687 |

Table S21: General linear model comparing **across male type** the **neuron/glia ratio of males (as residual of body size)** by **gonad weight of males (as residual of body size)** reveals no correlation in the morphs of *P. parae*, *P. picta* and *P. reticulata*. Below is the linear model and follow up comparison of slope (compared to 0 and pairwise comparisons) and intercept (pairwise comparisons). Significant effects in bold and underlined ($p<0.05$).

| Model Results (Neuron/Glia ratio ~ residual gonad weight*male type) | | | | |
|---|---|---|---|---|
| Effect | Estimate | Std. Error | t value | P value |
| **Linear Model** | | | | |
| **(Intercept)** | 0.53123 | 0.0389 | 13.6440 | 1.23e-16 |
| Residual Gonad Weight (ResGL) | 0.02010 | 0.2602 | 0.0772 | 9.39e-01 |
| Melanzona [*P. parae]* | 0.07650 | 0.0453 | 1.6884 | 9.91e-02 |
| Parae [*P. parae*] | -0.00773 | 0.0504 | -0.1534 | 8.79e-01 |
| *P. picta* | 0.13697 | 0.0451 | 3.0345 | 4.22e-03 |
| *P. reticulata* | 0.13389 | 0.0495 | 2.7052 | 9.98e-03 |
| ResGL: Melanzona [*P. parae]* | -0.60155 | 0.4688 | -1.2830 | 2.07e-01 |
| ResGL: Parae [*P. parae*] | -0.17530 | 0.3809 | -0.4602 | 6.48e-01 |
| ResGL: *P. picta* | -0.30168 | 0.3667 | -0.8227 | 4.16e-01 |
| ResGL: *P. reticulata* | 0.51401 | 0.4511 | 1.1395 | 2.61e-01 |
| **Slope to 0** | | | | |
| Immaculata [*P. parae*] | 0.0201 | 0.260 | 0.0772 | 0.939 |
| Melanzona [*P. parae*] | -0.5815 | 0.390 | -1.4909 | 0.144 |
| Parae [*P. parae*] | -0.1552 | 0.278 | -0.5579 | 0.580 |
| *P. picta* | -0.2816 | 0.258 | -1.0897 | 0.282 |
| *P. reticulata* | 0.5341 | 0.368 | 1.4496 | 0.155 |
| **Pairwise Slope Comparisons** | | | | |
| Immaculata [*P. parae*] – Melanzona [*P. parae]* | 0.602 | 0.469 | 1.283 | 0.703 |
| Immaculata [*P. parae*] – Parae [*P. parae*] | 0.175 | 0.381 | 0.460 | 0.990 |
| Immaculata [*P. parae*] – *P. picta* | 0.302 | 0.367 | 0.823 | 0.922 |
| Immaculata [*P. parae*] – *P. reticulata* | -0.514 | 0.451 | -1.140 | 0.785 |
| Melanzona [*P. parae*] – Parae [*P. parae*] | -0.426 | 0.479 | -0.890 | 0.899 |
| Melanzona [*P. parae*] – *P. picta* | -0.300 | 0.468 | -0.641 | 0.967 |
| Melanzona [*P. parae*] – *P. reticulata* | -1.116 | 0.537 | -2.079 | 0.249 |
| Parae [*P. parae*] – *P. picta* | 0.126 | 0.380 | 0.333 | 0.997 |
| Parae [*P. parae*] – *P. reticulata* | -0.689 | 0.462 | -1.493 | 0.573 |
| *P. picta* – *P. reticulata* | -0.816 | 0.450 | -1.813 | 0.381 |
| **Pairwise Intercept Comparisons** | | | | |
| Immaculata [*P. parae*] – Melanzona [*P. parae]* | -0.07650 | 0.0453 | -1.6884 | 0.45237 |
| Immaculata [*P. parae*] – Parae [*P. parae*] | 0.00773 | 0.0504 | 0.1534 | 0.99987 |
| **Immaculata [*P. parae*] – *P. picta*** | **-0.13697** | **0.0451** | **-3.0345** | **0.03245** |
| Immaculata [*P. parae*] – *P. reticulata* | -0.13389 | 0.0495 | -2.7052 | 0.07087 |
| Melanzona [*P. parae*] – Parae [*P. parae*] | 0.08423 | 0.0395 | 2.1311 | 0.22731 |
| Melanzona [*P. parae*] – *P. picta* | -0.06047 | 0.0325 | -1.8588 | 0.35563 |
| Melanzona [*P. parae*] – *P. reticulata* | -0.05739 | 0.0383 | -1.4966 | 0.57066 |
| **Parae [*P. parae*] – *P. picta*** | **-0.14470** | **0.0393** | **-3.6793** | **0.00587** |
| **Parae [*P. parae*] – *P. reticulata*** | **-0.14162** | **0.0443** | **-3.1998** | **0.02138** |
| *P. picta* – *P. reticulata* | 0.00308 | 0.0381 | 0.0807 | 0.99999 |
